# The histidine kinase NahK regulates pyocyanin production through the PQS system

**DOI:** 10.1101/2023.08.23.554518

**Authors:** Alicia G. Mendoza, Danielle Guercio, Marina K. Smiley, Gaurav K. Sharma, Jason M. Withorn, Natalie V. Hudson-Smith, Chika Ndukwe, Lars E. P. Dietrich, Elizabeth M. Boon

## Abstract

Many bacterial histidine kinases work in two-component systems that combine into larger multi-kinase networks. NahK is one of the kinases in the GacS Multi-Kinase Network (MKN), which is the MKN that controls biofilm regulation in the opportunistic pathogen *Pseudomonas aeruginosa (P. aeruginosa)*. This network has also been associated with regulating many virulence factors *P. aeruginosa* secretes to cause disease. However, the individual role of each kinase is unknown. In this study, we identify NahK as a novel regulator of the phenazine pyocyanin (PYO). Deletion of *nahK* led to a four-fold increase in PYO production, almost exclusively through upregulation of phenazine operon two (*phz2*). We determined that this upregulation is due to mis-regulation of all *P. aeruginosa* quorum sensing systems, with a large upregulation of the *Pseudomonas* quinolone signal (PQS) system and a decrease in production of the acyl-homoserine lactone-producing system, *las.* In addition, we see differences in expression of quorum sensing inhibitor proteins that align with these changes. Together, this data contributes to understanding how the GacS MKN modulates QS and virulence.

**Importance:** *Pseudomonas aeruginosa* is a Gram-negative bacterium that establishes biofilms as part of its pathogenicity. *P. aeruginosa* infections are associated with nosocomial infections. As the prevalence of multi-drug resistant *P. aeruginosa* increases, it is essential to understand underlying virulence molecular mechanisms. Histidine kinase NahK is one of several kinases in *P. aeruginosa* implicated in biofilm formation and dispersal. Previous work has shown that the nitric oxide sensor, NosP, triggers biofilm dispersal by inhibiting NahK. The data presented here demonstrates that NahK plays additional important roles in the *P. aeruginosa* lifestyle, including regulating bacterial communication mechanisms such as quorum sensing. These effects have larger implications in infection as they affect toxin production and virulence.

## Introduction

*Pseudomonas aeruginosa* is a Gram-negative bacterium that establishes biofilms as part of its pathogenicity. *P. aeruginosa* is associated with nosocomial infections that cause complications for patients with cystic fibrosis, cancer, or burn wounds (1). The most at-risk patient groups are those on ventilators, a problem intensified by increased ventilator use during the SARS-CoV-2 pandemic (2). Acute *P. aeruginosa* infections that lead to sepsis are typically associated with the planktonic state, where the bacteria are more motile and can travel around the body, infecting various organs rapidly (1, 3). Chronic *P. aeruginosa* infections can lead to pulmonary illnesses, especially in people who already have higher susceptibility to respiratory problems, such as cystic fibrosis patients (4). These chronic infections are usually associated with the biofilm or sessile state (1, 3). This planktonic-to-sessile switch is an essential part of *P. aeruginosa* virulence.

The GacS Multi-Kinase Network (MKN) regulates the planktonic-to-sessile switch (3, 5). Several kinases, including GacS, PA1611 (PA14_43670), RetS, SagS, and NahK, signal in this network to regulate the activity of a post-transcriptional global regulator protein, RsmA (3, 5). RsmA inhibits translation of mRNAs related to biofilm formation, quorum sensing (QS), pyocyanin (PYO) production, and type VI secretion systems, promoting the planktonic state (3, 5). RsmA is inactivated by the small regulatory RNAs, *rsmY* and *rsmZ,* which are transcribed in response to MKN activity (3, 5). All kinases in this network are hybrid histidine kinases that respond to extracellular signals which include nitric oxide (NO), glycan mucins and calcium ions. However, further study is needed to identify additional signals and determine the physiological consequences for how these signals modulate RsmA activity (5, 6, 7, 8).

NO is a diatomic gas that signals biofilm dispersal in many bacteria at low nanomolar to picomolar concentrations (9). In *P. aeruginosa*, the NO sensing protein NosP is necessary for biofilm dispersal (6). At a molecular level, NosP functions by inhibiting its cocistronic-associated kinase, NahK, when in the ferrous NO-ligated state (6). NahK is one of four kinases that regulate the phosphorylation state of HptB, which is one of the main response regulators in the GacS MKN (6, 11). When HptB is not phosphorylated, HptB indirectly activates transcription of *rsmY,* leading to inactivation of RsmA and, thus, promotes biofilm formation (5). NosP and NahK have also been identified as part of the *Pseudomonas* biofilm transcriptome through comparative transcriptome analysis of 138 biofilm-forming clinical *Pseudomonas aeruginosa* isolates (11). Outside of these works, not much is known about the molecular roles of NosP and NahK in biofilm regulation and other cellular processes, even though the GacS MKN is implicated in essential bacterial processes including QS, antibiotic resistance, metabolism, replication, and virulence (3, 11, 12).

In *P. aeruginosa*, three QS systems act sequentially, *las*, *rhl* and *pqs* (12). RsmA and the GacS MKN control QS and phenazine production as part of their global regulon (5). When RsmA is active, it inhibits translation of *lasR* and *rhlR* (12). At high cell density, RsmA repression is relieved, allowing for QS to occur, which also results in phenazine production (12). PYO production is controlled, in part, by all three QS systems that contribute to the transcriptional activation of two phenazine-producing operons, *phzA1-G1* (*phz1*) and *phzA2-G2* (*phz2*) (12, 13, 14, 15). These operons are ∼98% identical at the genomic level and each one is sufficient to confer production of phenazine-1-carboxylic acid (PCA), a precursor to the well-characterized blue phenazine, PYO. LasR and RhlR are known to activate *phz1* directly, while the PQS influence on *phz2* is indirect through an unknown mediator (16). These operons are also controlled by additional factors, such as other transcription factors and QS inhibitor proteins (12, 17). While both operons contribute to phenazine production, *phz2* is more important than *phz1* for host colonization in mouse models (13). Here, we describe a novel role for NahK in modulating of the PQS system, which results in downregulation of *phz2* expression, and, therefore, also decreases PYO production.

## Results

### Deletion of nahK leads to overproduction of the phenazine PYO

After discovering the role of NosP and NahK in biofilm dispersal, we became interested in how these proteins individually contribute to RsmA-dependent phenotypes (6). We generated a genetic deletion of *nahK* and found that this strain secreted a blue pigment into the culture supernatant (**Figure 1A**). The blue coloration indicated the phenazine PYO, which we verified using acidified-chloroform extraction of supernatant and UV-visible light spectroscopy. (**Figure 1B**) (14). The Δ*nahK* strain showed a four-fold increase of PYO in planktonic culture and a two-fold increase in biofilms (**Figure 1C and 1D**) (18). PYO production was reduced by complementation of *nahK* in both planktonic and biofilm cultures (**Figure 1C and 1D**).

**Figure 1.**
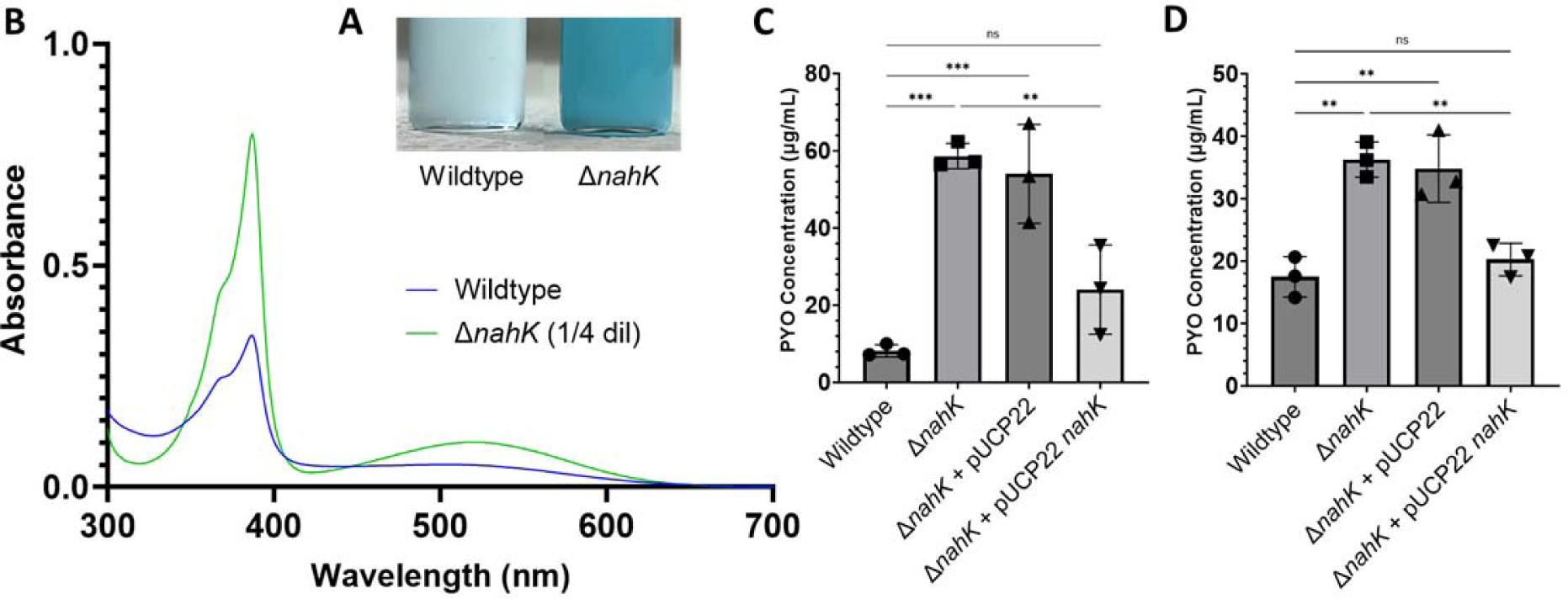
Pyocyanin is overproduced in Δ*nahK*. **(A)** Chloroform extract of Δ*nahK* supernatant has a bright blue color. **(B)** PYO is characterized by a peak at 520nm in 0.1M HCl (18). Δ*nahK* is diluted 4-fold in 0.1M HCl compared to wildtype. **(C, D)** Beer’s law quantification of pyocyanin production at 520nm (extinction coefficient 17.072 from 18). (**C**) Planktonic, liquid cultures (n = 3) and **(D)** biofilm, agar cultures. n = 3. *p* values were calculated using one-way ANOVA and a Tukey multiple comparisons test. * p ≤ 0.5.

Many factors in the GacS MKN have been shown to influence PYO production (5, 8, 19, 20). Deletion of *hptB* leads to decreases in motility and virulence, but effects on PYO or other toxins were not reported (10, 21). *gacS* and *gacA* deletions have shown decreases in virulence and biofilm production, with some reports showing a two-fold decrease in PYO production (8, 20, 22). In PAO1, Δ*rsmY* and Δ*rsmZ* have been shown to have decreases in PYO production, while Δ*rsmA* has a modest increase in PYO levels, about two-fold compared to wildtype (20). Recently, Δ*retS* in PAO1 has been shown to also have an increase in PYO production similar to Δ*rsmA*, with an increase about two-fold (8). Overall, this suggests a novel regulation of PYO by NahK in this network.

### ΔnahK is more virulent than PA14 wildtype

PYO is a redox-active pigment that contributes to *P. aeruginosa* pathogenicity by generating reactive oxygen species. For example, PYO has been shown to result in death of neutrophils by interfering with their mitochondrial respiratory chain (23). To determine if Δ*nahK* is more virulent than wildtype *P. aeruginosa* PA14, we performed a *Caenorhabditis elegans* slow killing assay. Over four days, only 36% of worms survived that were fed with Δ*nahK*, compared to 78% survival of worms fed with wildtype (**Figure 2**). This reduction was partially restored by complementing the Δ*nahK* strain with *nahK* (**Figure 2**). We have also shown a similar phenotype in a mung bean virulence model that demonstrated Δ*nahK* kills sprouts more readily than wildtype (Supp Figure 1).

**Figure 2.**
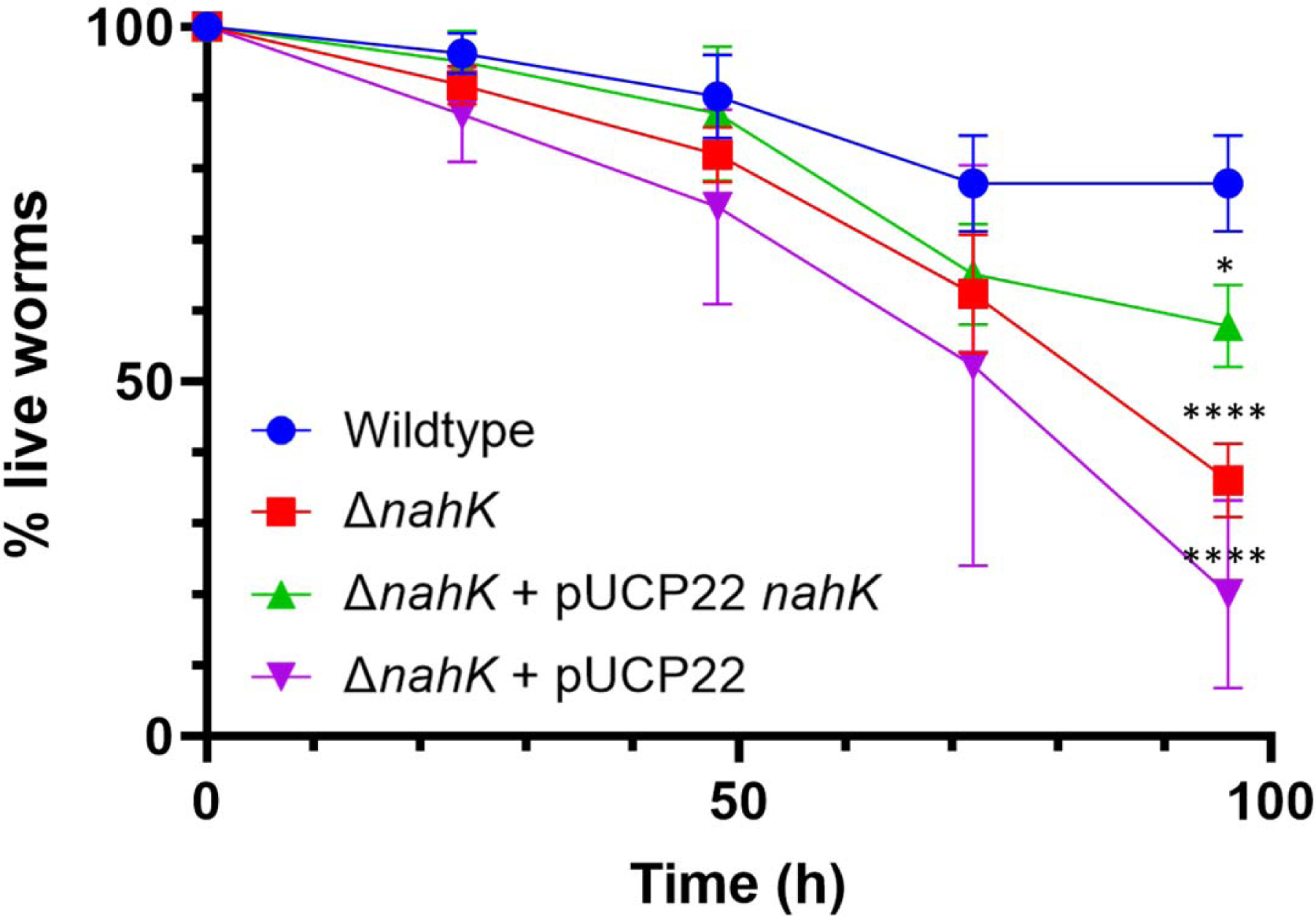
Δ*nahK* is more virulent than PA14 wildtype. Slow-killing kinetics of all strains in the nematode *C. elegans*. After 5 days of exposure to the bacteria, wildtype shows ∼20% killing, while ΔnahK shows ∼55% killing. Error bars represent the standard deviation of at least 4 biological replicates, with each replicate consisting of 30-35 worms. *p* values were calculated using one-way ANOVA. * p ≤ 0.5, **** p ≤ 0.0001.

### ΔnahK mis-regulates phenazine biosynthesis genes

We then set to determine the molecular mechanism of NahK-dependent regulation of PYO production. Phenazine biosynthesis occurs through a branched, multi-step pathway (24). The first step is the conversion of chorismic acid to phenazine-1-carboxylic acid (PCA) by the two redundant phenazine operons, *phz1* and *phz2*, which have ∼98% DNA sequence homology (13, 16, 24). Despite their conservation, the two operons are under the control of different promoter sequences, leading to differential regulation (13, 16). In PA14, both *phz1* and *phz2* contribute to phenazine production in planktonic cultures while, during biofilm growth, *phz2* is the dominant operon (13). Therefore, we investigated which *phz* operon was responsible for NahK-dependent PYO production in planktonic culture.

To study the transcriptional activity of each operon, we generated mScarlet transcriptional reporters with the promoters of *phz1* or *phz2* driving mScarlet expression and genomically integrated them into wildtype and Δ*nahK* strains. We then grew the strain planktonically for 24 hours and tracked the mScarlet fluorescence signal (**Figure 3A**). We found that expression of the P*phz2-*mS reporter was increased in both strains compared to P*phz1-*mS. P*phz1-*mS had minimal expression in both wildtype and Δ*nahK* (**Figure 3A**). Δ*nahK*::P*phz2*-mS had higher expression than the wildtype, suggesting that *phz2* was upregulated in Δ*nahK*, giving rise to the PYO overproduction phenotype (**Figures 3A and 1**). This same trend was also observed in biofilm; however, the differences were not statistically significant (**Figure 3B**). This trend is expected since phenazine production in biofilms is mostly *phz2*-dependent (13).

**Figure 3.**
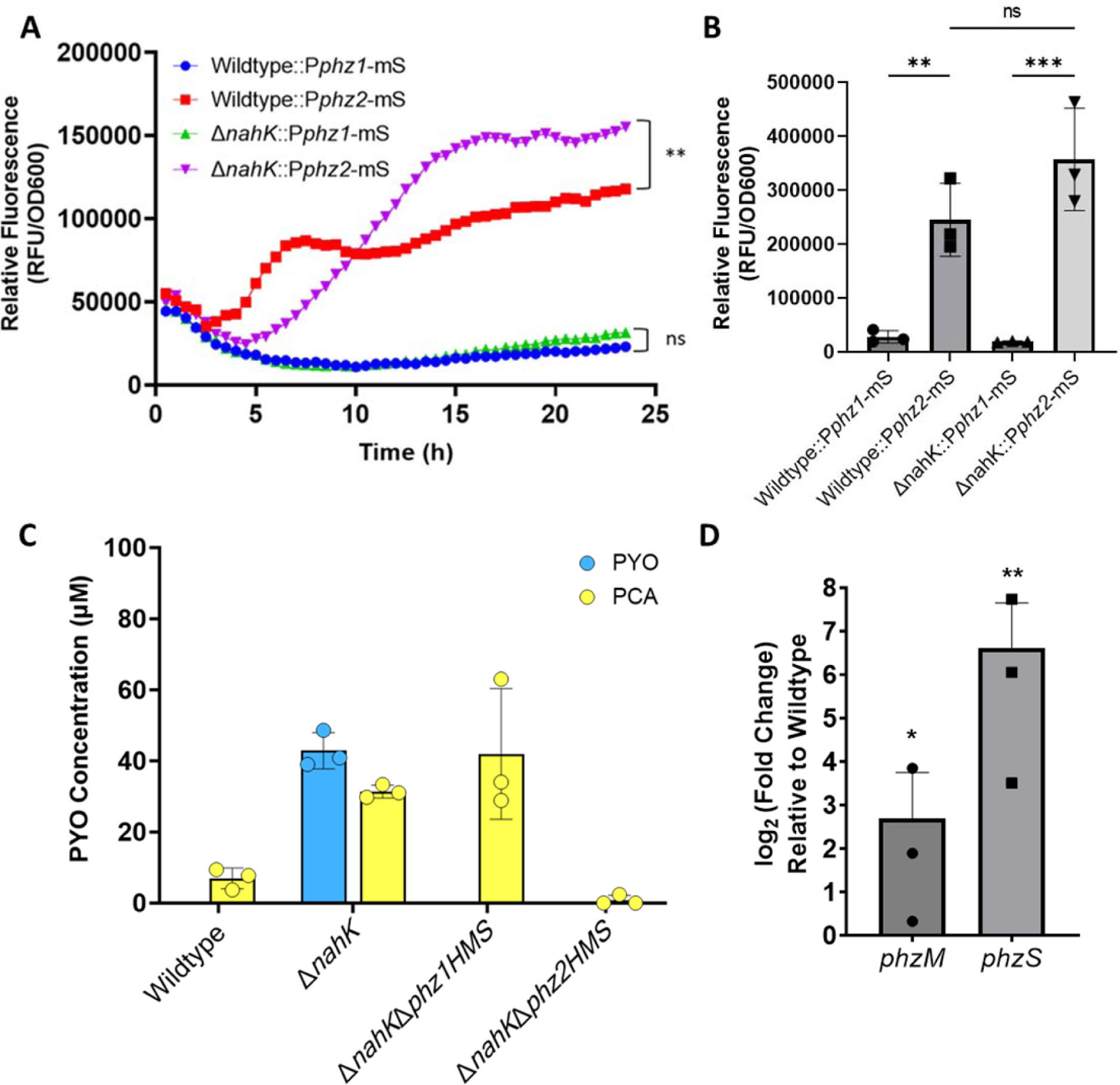
*Phz2* is the driver of PYO production. (**A**) P*phz2-*mS is expressed more than P*phz1-*mS in the wildtype and Δ*nahK* planktonically. Δ*nahK* has higher expression of P*phz2-*mS than wildtype overall. *p* values were calculated using one-way ANOVA and a Tukey multiple comparisons test at final time point. n = 3. (**B**) P*phz2-*mS is expressed more than P*phz1-*mS in the wildtype and Δ*nahK* biofilms. *p* values were calculated using one-way ANOVA and a Tukey multiple comparisons test. n = 3. (**C**) HPLC quantification of phenazines PCA and PYO in wildtype, *ΔnahK, ΔnahKΔphz1HMS* and *ΔnahKΔphz2HMS.* n=3 (**D**) qPCR for *phzM* and *phzS* in *ΔnahK* compared to wildtype. Gyrase A (*gyrA*) was used as a housekeeping gene (31). *p* values were calculated using unpaired, two-tailed *t-*tests comparing ΔnahK ΔCt values to wildtype ΔCt values. n = 3. * p value < 0.05 ** p value < 0.01 *** p value < 0.001.

The conversion of PCA to PYO is mediated by two enzymes: the SAM-dependent methyltransferase PhzM converts PCA to 5-methylphenazine-1-carboxylic acid betaine (24, 25). The oxygen-dependent monooxygenase PhzS converts 5-methylphenazine-1-carboxylic acid betaine to PYO (26, 27, 28). PhzS can also use PCA as a substrate to make 1-hydroxyphenazine (24, 29, 30). To further determine that *phz2* drives PYO production in Δ*nahK*, we generated Δ*nahK* strains that contained deletions of either *phz1* or *phz2* and quantified phenazine production. These strains were also Δ*phzH*Δ*phzM*Δ*phzS* (ΔHMS), therefore the only phenazine produced is PCA. When phenazines were quantified, Δ*nahK*Δ*phz1HMS* produced PCA levels comparable to Δ*nahK* (**Figure 3C**). Δ*nahK*Δ*phz2HMS* generated nearly no detectable PCA, suggesting that nearly all phenazines produced by Δ*nahK* are produced by *phz2* (**Figure 3C**).

To assess if PhzM and PhzS are affected by NahK, we assessed the expression of these corresponding genes in Δ*nahK* (**Figure 3D**). Indeed, both *phzM* and *phzS* were upregulated in Δ*nahK* compared to wildtype (**Figure 3D**).

### Quorum sensing is mis-regulated in ΔnahK

Both *phz* operons, *phzM* and *phzS* are regulated by quorum sensing (QS) (16). To understand how QS systems are involved in modulating PYO production in our system, we performed qPCR on the regulators of the *phz* operons (**Figure 4**). *Phz1* is activated by two of the main QS pathways, the *las* and *rhl* systems (12). LasI and RhlI are lactone synthases that make N-acyl homoserine lactones, which activate the transcription factors LasR and RhlR (16, 32, 33). In agreement with **Figure 3**, factors that activate *phz1* showed no difference in transcriptional levels in *ΔnahK* compared to wildtype levels (**Figure 4**). *phzM* and *phz*S are also thought to be controlled primarily through the *las* and *rhl* systems because these genes flank the *phz1* operon. This may suggest a role for the *las* and *rhl* systems in the *ΔnahK* phenazine phenotype that is not appreciated by the fluorescent transcriptional reporter.

**Figure 4.**
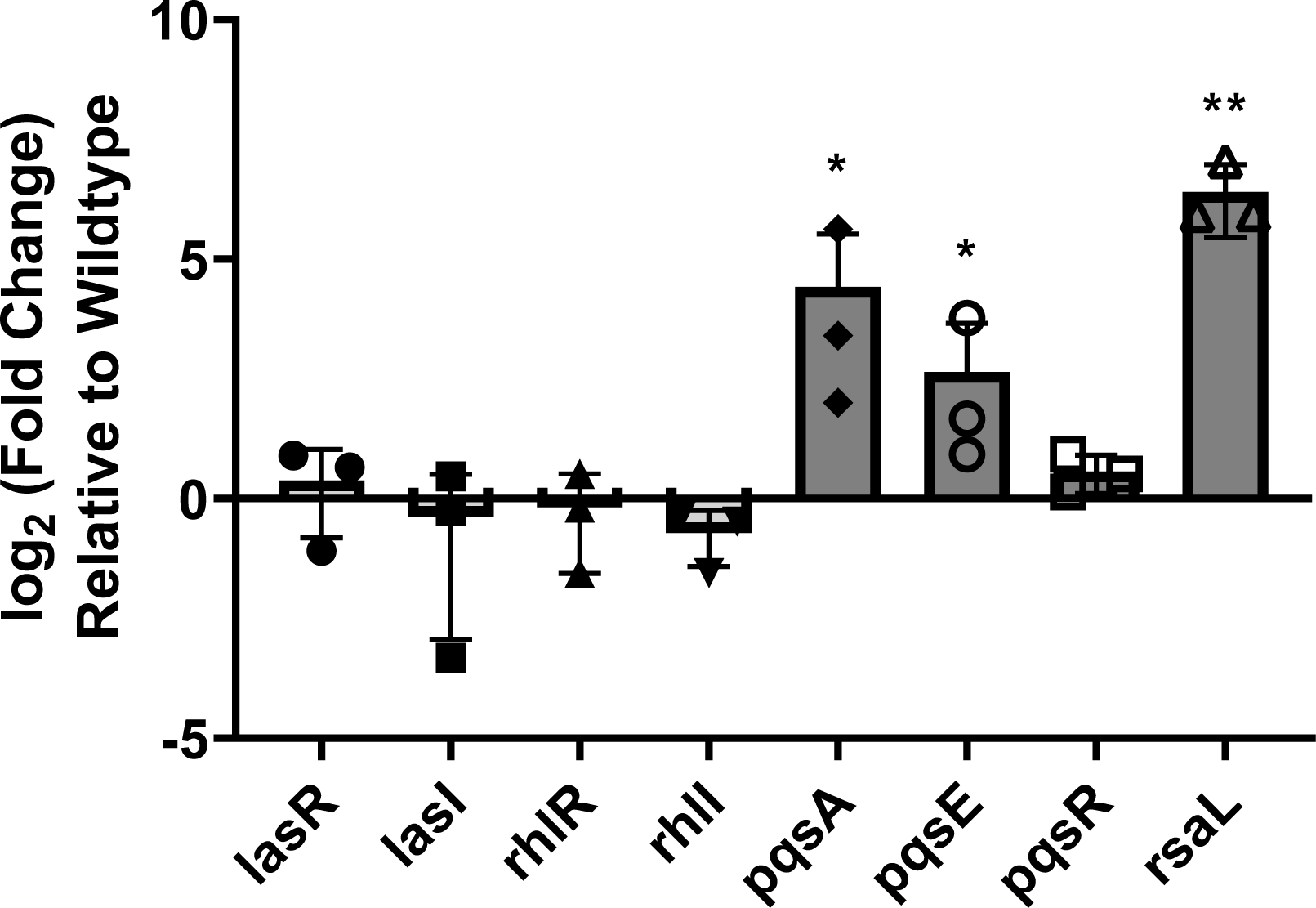
Pyocyanin production may be regulated by the PQS system and *rsaL*. qPCR for the main quorum sensing transcriptional regulators in *ΔnahK* compared to wildtype. Pyocyanin production is potentially induced by each of these transcriptional regulators. Gyrase A (*gyrA*) was used as a housekeeping gene (31). Bars above and below the threshold represent up- and downregulation, respectively. *p* values were calculated using unpaired, two-tailed *t-*tests comparing ΔnahK ΔCt values to wildtype ΔCt values. n = 3. * p value < 0.05 ** p value < 0.01.

Regulation of *phz2* is not mediated by LasR/RhlR directly and is overall not as well understood as that for *phz1* (16, 13). *phz2* is activated primarily by two factors: QS-inhibitor protein RsaL and *Pseudomonas* quinolone signal, PQS (13). RsaL is a global transcription factor that can be activated by LasRI and is involved mostly in combating oxidative stress (16). It is also mostly studied within the context of the *las* system, as RsaL represses the *las* operon (34). It is believed that RsaL indirectly controls *phz2* via an unidentified regulator (16).

*Pseudomonas* quinolone signal (PQS) is a QS molecule that has additional functions of iron binding and antioxidant properties (35). *phz2* expression correlates directly with PQS production; when PQS is upregulated, *phz2*-mediated PCA production is upregulated (16). PQS is also synthesized from precursor molecule chorismic acid and converted to PQS by the *pqs* operon which includes *pqsABCD* and *pqsE* (35). Current research suggests *pqsE* can be differentially regulated in the absence of RhlR and compensate for some RhlR-mediated transcriptional regulation (36). MvfR/PqsR senses PQS and regulates the *pqs* operon (12). Genes in the *pqs* operon and *rsaL* are highly upregulated in *ΔnahK* compared to wildtype (**Figure 4**).

To corroborate these results, we performed untargeted-LCMS to quantify QS molecules from the bacterial supernatant of the wildtype and *ΔnahK* strains (**Figure 5**). PQS derivatives, like 2-heptyl-4-quinolone (HHQ) and dihydroxyquinoline (DHQ) were more prevalent in *ΔnahK* compared to wildtype (**Figure 5**). Interestingly, we found a reduction in N-(3-oxododecanoyl)-L-homoserine lactone (C12-HSL) production in *ΔnahK,* suggesting that there may be post-transcriptional regulation of the *las* quorum sensing system. Many pathogenic and clinically relevant strains of *Pseudomonas aeruginosa* have defects or deletions in the *las* system (37). This may relate to other ways *ΔnahK* is more virulent than wildtype, possibly separate from PYO production (**Figure 2**). RsmA activity is also known to inhibit the Las system post-transcriptionally, which overall promotes pathogenicity (38). Because NahK activity may regulate RsmA activity through the HptB branch of the GacS MKN, this may suggest that deletion of *nahK* improperly promotes RsmA activity, leading to these changes in QS.

**Figure 5.**
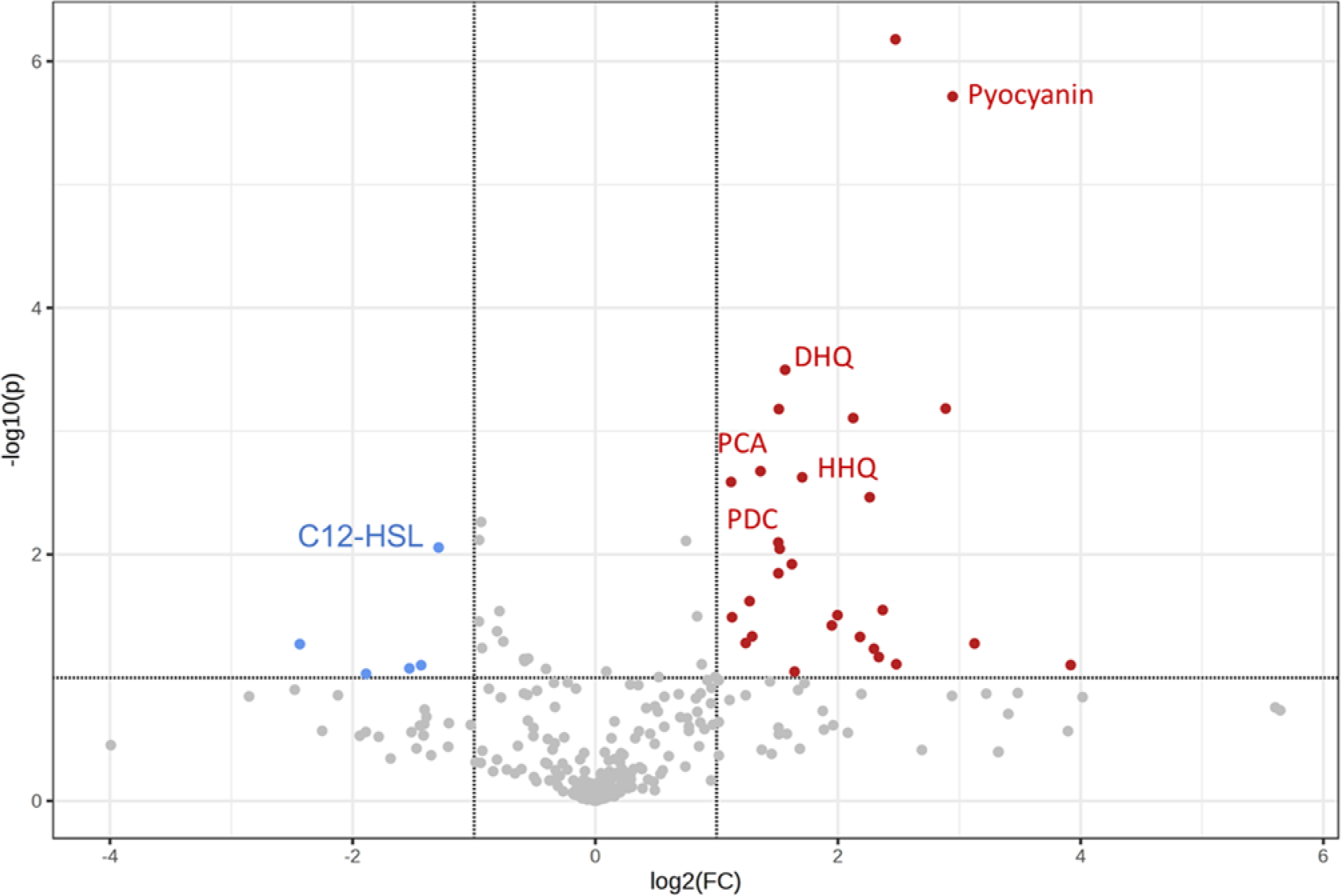
PQS precursors are overexpressed in *ΔnahK* supernatant, while C12-HSL is reduced. Values are a ratio of prevalence in Δ*nahK*:wildtype in positive mode LCMS. Bars above and below the threshold represent up- and downregulation, respectively. *p* values were calculated using unpaired, two-tailed *t-*tests unpaired, two-tailed *t-*tests (Supplemental Table 4).

### Quorum sensing is mis-regulated at both a transcriptional and post-translational level

QS systems in *P. aeruginosa* have several layers of regulation. First, these systems work sequentially as a function of cell density and regulate each other (the *las* system activates the *rhl* system, which in turn inactivates *las*). A similar feedback mechanism has been described for *rhl* and *pqs,* where RhlR activates the *pqs* operon and PqsR turns off the *rhl* operon (12). QS is also regulated by QS inhibitor proteins, many of which need further characterization (12). The best-characterized inhibitors include RsaL and QscR, inhibitors of the *las* system, QteE, inhibitor of the *rhl* system, and QslA, inhibitor of the *pqs* system (39, 40, 41). The mechanism of inhibition is typically by protein-protein interaction, where the inhibitor binds the transcription factor, inactivating it (12). Because these are protein-protein interactions, we hypothesized that if inhibition of the *las* and *rhl* system were occurring, this inhibition would not be reflected in the transcription levels of the transcription factors LasR and RhlR via qPCR (**Figure 4**). Using qPCR, we examined relative levels of each inhibitor in Δ*nahK* compared to wildtype (**Figure 6**). As expected, we saw an increase in QS inhibitors for the *las* and *rhl* systems, and a decrease in the inhibitor for the *pqs* system (**Figure 4**, **Figure 6**).

**Figure 6.**
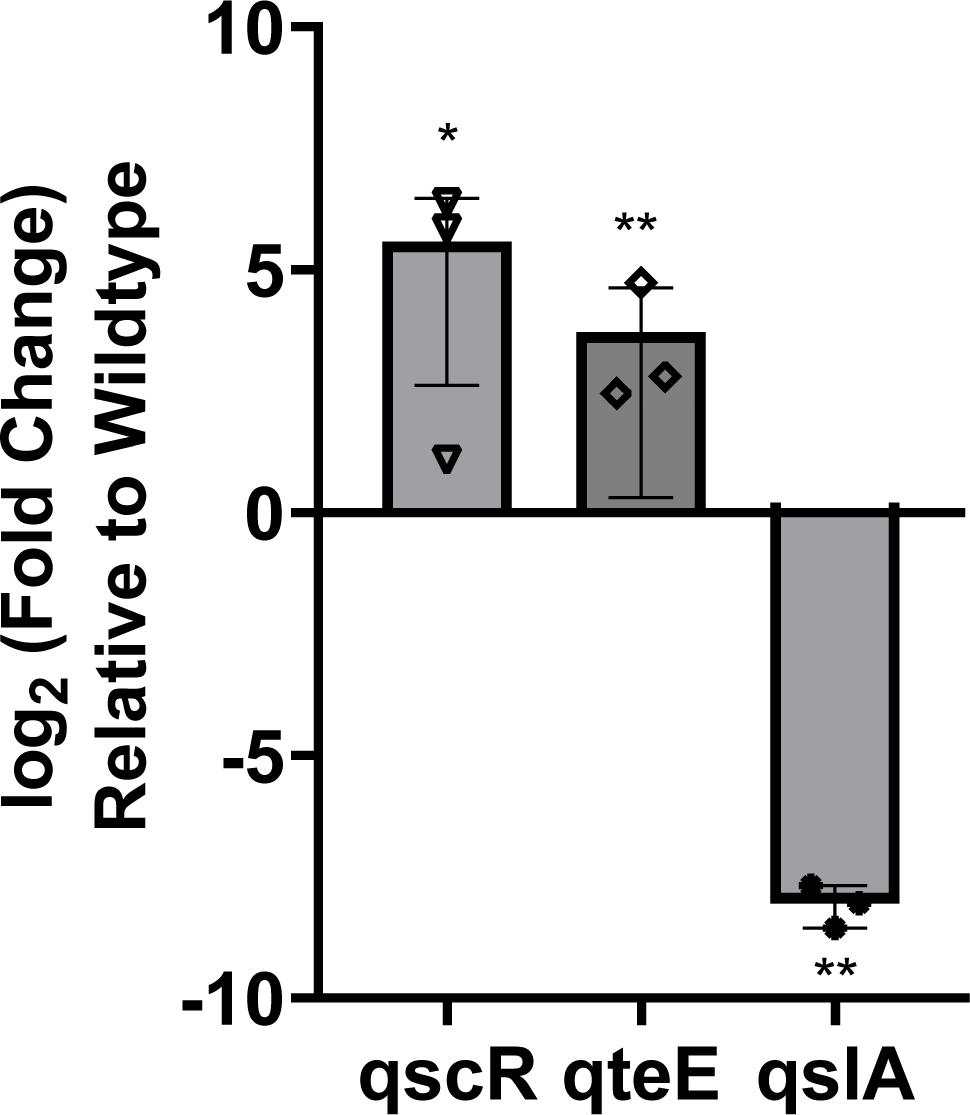
Quorum sensing inhibitors for the *las* and *rhl* systems are upregulated, whereas the *pqs* inhibitor is downregulated. qPCR for the main QS inhibitors in *ΔnahK* compared to wildtype. Gyrase A (*gyrA*) was used as a housekeeping gene. Bars above and below the threshold represent up- and downregulation, respectively. *p* values were calculated using unpaired, two-tailed *t-*tests comparing ΔnahK ΔCt values to wildtype ΔCt values. n = 3. * p value < 0.05 ** p value < 0.01.

### PQS promotes phz2 expression in ΔnahK

Once we found PQS was increased in Δ*nahK*, we then asked if PQS was increasing *phz2*-mediated PYO production in our strains. To determine this, we devised a co-culture experiment. A donor strain, either wildtype or Δ*nahK*, would provide PQS to a recipient strain that could not produce PQS (Δ*pqsABC*) but did encode for either the *phz1* or *phz2* transcriptional mScarlet reporter (Δ*pqsABC*::P*phz1*-mS and Δ*pqsABC*::P*phz2*-mS). Upon co-culturing, we could track activation of the *phz* reporters as a function of growth. Wildtype or Δ*nahK* would provide PQS that activates *phz2* in Δ*pqsABC*::P*phz2*-mS in a manner dependent on amount of PQS secreted. Δ*pqsABC*::P*phz-*mS reporter strains did not activate the reporters during their own growth, only when co-cultured (**Supp. Figure 2**). When co-cultured with wildtype, *phz2* activated more than *phz1*, similar to our previous finding in **Figure 4** (**Figure 7A**). When co-cultured with Δ*nahK*, *phz2* activated more than *phz1*, and *phz2* activation was increased compared to wildtype *phz2* activation levels (**Figure 7A**). To further confirm this was due to PQS itself, we also generated Δ*pqsR*::P*phz* reporter recipient strains that are unable to respond to PQS. These strains, when co-cultured with either wildtype or Δ*nahK*, showed no activation of either reporter (**Supp. Figure 3A**). In addition, co-cultured wildtype or Δ*nahK* with Δ*pqsABC*::P*phz2*-mS showed increased activation in a 3-day colony biofilm (**Figure 7B**).

**Figure 7.**
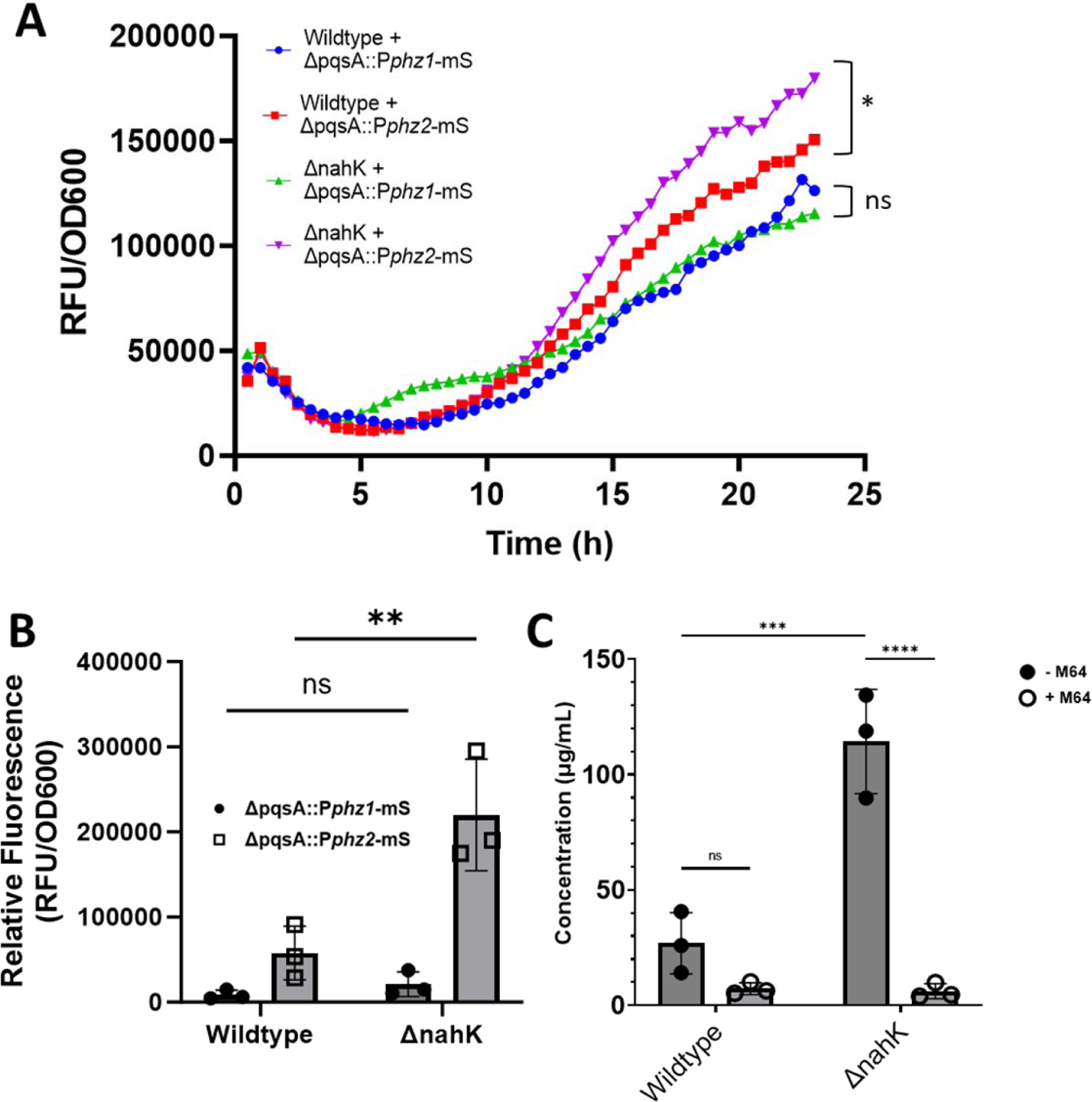
PQS drives *phz2* in ΔnahK. (**A**) Co-cultured wildtype or Δ*nahK* with Δ*pqsABC* (planktonic) containing either the *phz1* or *phz2* mScarlet reporter shows increased activation of *phz2* with both donors, with a higher activation in Δ*nahK*. n = 3. *p* values were calculated using one-way ANOVA and a Tukey multiple comparisons test at final time point. * p ≤ 0.05 (**B**) 3-day biofilms of co-cultured wildtype or Δ*nahK* with Δ*pqsABC* containing either the *phz1* or *phz2* mScarlet reporter shows increased activation of *phz2* with both donors, with a higher activation in Δ*nahK*. n = 3. *p* values were calculated using one-way ANOVA and a Tukey multiple comparisons test at final time point. * p ≤ 0.05 (**C**) Beer’s law quantification of PYO production at 520nm in planktonic, liquid LB cultures with and without exposure to 1µM of M64 PqsR inhibitor (42). *p* values were calculated using one-way ANOVA and a Tukey multiple comparisons test. *** p ≤ 0.001, **** p ≤ 0.0001.

We then determined PYO concentrations in wildtype and Δ*nahK* supernatants after exposure to 1µM of the PqsR inhibitor, M64 (42). With M64, both wildtype and Δ*nahK* had diminished PYO production, further suggesting PQS was driving PYO production (**Figure 7C**). Overall, this suggests that ΔnahK overexpresses *phz2*-mediated PYO because the strain generates an increased amount of PQS.

## Discussion

Here, we describe the effect of the histidine kinase (HK) NahK on the PQS-dependent production of the phenazine PYO in *P. aeruginosa* PA14 (**Figure 8**). NahK is conserved in many Gram-negative bacteria, and NO regulation of QS systems has been studied in *Vibrio cholerae, Vibrio harveyi* and *Staphylococcus aureus* (43, 44, 45). In *V. cholerae*, the *nosP/nahK* operon encodes for *Vc*NosP, a NO sensor, and hybrid HK VpsS. The mechanism is similar; NO-bound NosP (VpsV) inhibits VpsS, leading to dispersal. There, VpsS affects QS by transferring a phosphoryl group to LuxU, and subsequently to the transcription factor LuxO, which results in increased virulence and biofilm at low cell density (43). Therefore, it was reasonable to hypothesize that in *P. aeruginosa* the *nosP/nahK* operon would influence QS and its downstream effectors. However, it is surprising how large of an effect Δ*nahK* has on PYO production that was previously unexplored.

**Figure 8.**
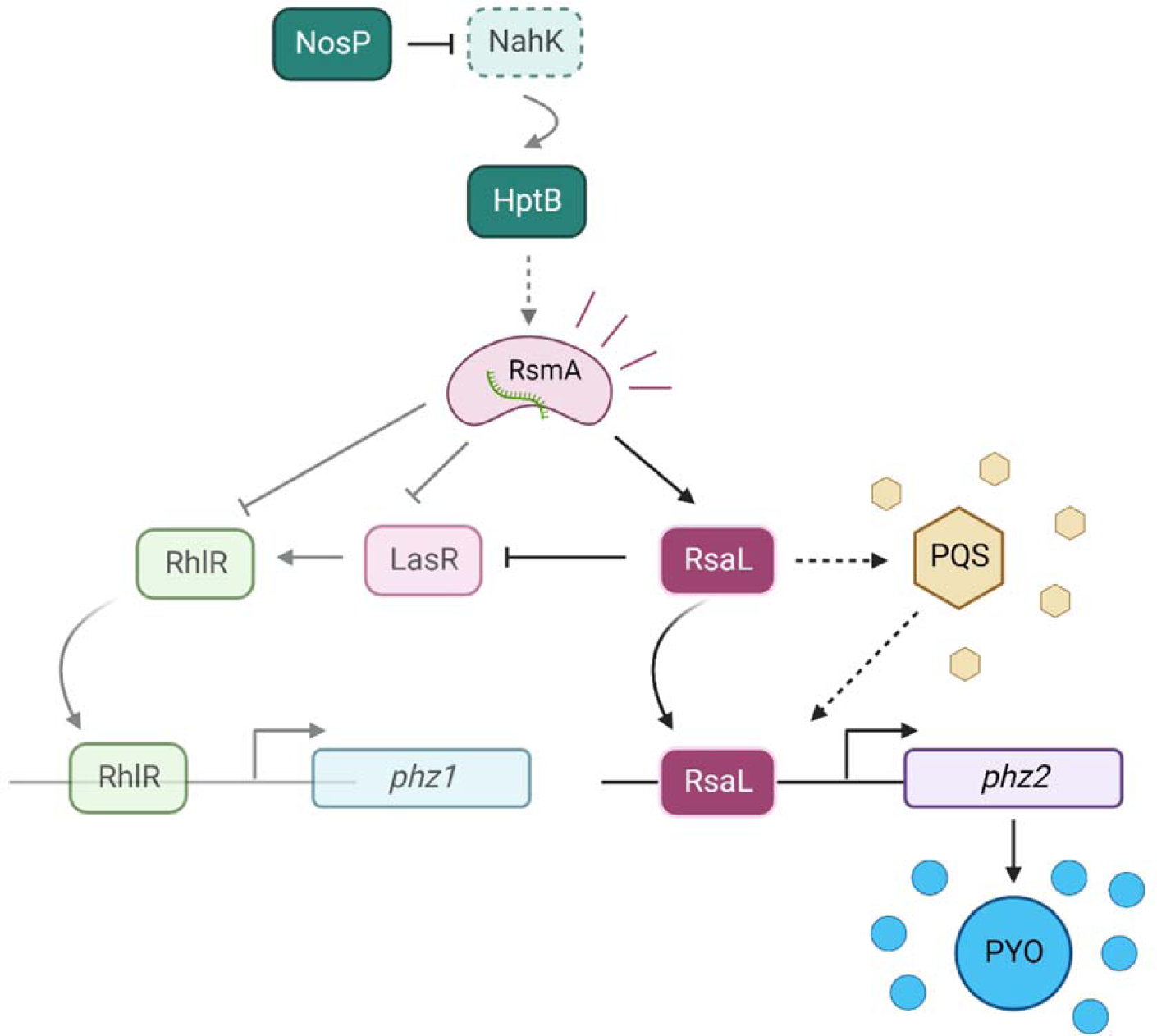
Graphical abstract. Inhibition of NahK leads to increased RsmA signaling. This signaling represses *las* and *rhl* systems but activates PQS signaling. These fluctuations in signaling lead to an upregulation of *phz2*-mediated PYO production.

The GacS MKN that NahK is implicated in has been shown to indirectly affect PYO production through RsmA regulation of QS (3, 5). In PAO1, Δ*rsmY* and Δ*rsmZ* have shown to decrease PYO, while Δ*rsmA,* Δ*gacS* and Δ*retS* have shown increases in PYO (8, 20, 46). Many studies focus on how these GacS MKN factors influence the Las and Rhl systems, suggesting these modulations of PYO are potentially through a *phz1-*mediated overexpression, rather than a *phz2*-mediated one (5). Our data instead suggest these changes in PYO may be the result of changes in PQS production and *phz2* expression. However, deletions of all factors in the GacS MKN need to be characterized more robustly in PA14 in order to establish this mechanism.

The data presented here could also suggest that there is a more direct link between NahK and QS. It is possible that NahK can either heterodimerize with other kinases in *P. aeruginosa* or phosphorylate other proteins. Recently, these types of interactions have been appreciated for other kinases in the GacS MKN. Orphan sensor kinase SagS is known to crosstalk outside of the GacS MKN with BfiS and NicD to regulate cyclic-di-GMP and *rsmZ* levels, both to control biofilm formation in a Gac-independent manner (19). In *Shewanella oneidensis*, *So*NahK is known to phosphorelay with three response regulators, leading to transcriptional and cyclic-di-GMP production changes (47). Based on what is known in *S. oneidensis* and other kinases in the *P. aeruginosa* GacS MKN, it is possible that NahK could have another unidentified response regulator. This requires additional research.

It is also surprising that *nahK* has not appeared in screens for regulators of PYO. It has been suggested that the *nosP/nahK* operon is QS-regulated (11, 48, 49). Letizia *et. al.* suggests that, in the absence of all other QS transcription factors and signaling molecules, RhlR downregulates the *nosP/nahK* operon in PAO1 (48). Perhaps NahK functions as an important intermediate to transition from the Las/Rhl systems to the PQS system during stationary phase. There may be a mechanism where RhlR transcriptionally represses NahK to lower NahK signaling, which would therefore activate the PQS system, similar to how deletion of *nahK* promotes PQS production as shown here. However, additional research is required to know when NosP and NahK are present and active in the bacteria.

NahK may also play an important role in connecting PYO, biofilm regulation and anaerobic respiration. Intracellular NO levels in *P. aeruginosa* are indirectly regulated by all three QS systems because they play a role in controlling the denitrification machinery (50). PYO is thought to be used as an extracellular electron shuttle to promote anaerobic respiration in deep layers of biofilms that are not exposed to the environment (51, 52). It is possible that NO produced as a byproduct of anaerobic respiration is activating NosP signaling, which therefore inhibits NahK activity and through modulating QS systems, produces more PYO to promote more anaerobic respiration. Because of this, it is possible NahK has not shown up in previous screens for phenazine regulators since its main role is specific to microaerobic conditions.

Overall, understanding how NahK links NO and QS may have important implications for how NO sensing controls molecular mechanisms in the bacteria. In infection, where immune cells secrete NO to combat bacterial invasion, it may be advantageous for NO to trigger cell density independent PQS signaling through NosP inhibition of NahK. The PQS system drives activation of cytotoxicity through phenazine production, as well as activates iron acquisition systems and modulates host immune signaling (35). Therefore, NO signaling may be one early way bacteria activate counter-mechanisms to survive in a host because NosP responds to signaling levels of NO (6). Further characterization of the physiological relevance of NosP-mediated NO signaling during infection is necessary to understand how NO and QS integrate environmental signals.

## Materials and Methods

### Bacterial strains, growth conditions and media

The bacterial strains and plasmids used in this study are described in **Supplemental Table 1 and 2**. Oligonucleotides are described in **Supplemental Table 3**. *P. aeruginosa* strains were grown aerobically in Luria-Bertani (LB) broth or succinate-based minimal media (35mM K2HPO4, 22mM KH2PO4, 7.6 mM (NH4)Cl, 1.7mM MgSO4, 40mM succinic acid, and 27.5mM NaOH, pH 7.0) at 37°C (53). *E. coli* strains were grown aerobically in LB broth or on LB agar at 37°C.

### Generation of deletion strains

Markerless deletions were generated using methods previously described (8). In brief, 1 kb of flanking sequence for target locus was amplified and inserted in pMQ30 using gap repair cloning into *Saccharomyces cerevisiae* InvSc1. Plasmids were than transformed in E. coli donor strain WM3064 and conjugated into *P. aeruginosa* PA14 and selected for on LB agar plates containing 100 µg/mL gentamicin. Clones were confirmed via PCR.

### Pyocyanin extraction

PYO extraction was performed as described previously elsewhere (14, 54, 55). In brief, for planktonic culture, supernatant was collected from 100mL succinate media cultures grown for 16-18hours. PYO was extracted via chloroform extraction, then extracted into 0.1M HCl for quantification. UV visible light spectra was taken on a Varian Cary 100 Bio Spectrophotometer. PYO was quantified using Beer’s law (extinction coefficient 17.072 µg/mL) (54). For biofilm extraction, succinate-based minimal media agar plates (1% agar), were prepared using 35mmx10mm petri dishes. Bacteria cultures were adjusted to an OD of 0.8 before spotting onto agar plate. Biofilms were grown for three days at 25°C in the dark. The biofilm and agar were harvested and submerged in 4mL of chloroform overnight 25°C in the dark. PYO was quantified as described above. For PYO quantification with exposure to M64 PqsR inhibitor, LB cultures at OD_600_ of 0.05 were exposed to 1µM of M64 and grown for 16-18 hours. PYO was extracted and quantified as described above.

### *C. elegans* Slow Killing Assay

*C. elegans* slow killing assay has been previously described as an effective method to observe *P. aeruginosa* virulence (51). 100 μL of PA14 wildtype and PA14 *ΔnahK* were spotted onto slow killing agar plates (0.3% NaCl, 0.35% Bacto-Peptone, 1 mM CaCl2, 1 mM MgSO4, 5 µg/ml cholesterol, 25 mM KPO4, 50 µg/ml FUDR, 1.7% agar). Plates were subsequently incubated 24 hours at 37°C and then left for 48 hours at room temperature. 30-35 larval stage 4 *C. elegans* were transferred onto PA14 WT and PA14 *ΔnahK* seeded plates. Live worms were counted for four days.

### Mung Bean Virulence Assay

To assess the comparative virulence of PA14 Δ*nahK* to PA14 wildtype, mung beans were exposed as described in Garge et al. with some minor modifications (56). Briefly, mung beans (Cool Beans N Sprouts) were washed in 70% (v/v) EtOH/Water, a solution of 30% commercial bleach and 0.02% Triton X-100 and then rinsed 3x with sterile ddH_2_O. Sterilized mung beans were placed onto water agar (0.8%) and supplemented with 2 mL of sterilized ddH_2_O and wrapped to maintain. Seeds were left to germinate for 24hrs at 37C. After germinating, sprouts of similar length and appearance were selected and randomly sorted into groups. Prepared overnight cultures of PA14 WT and PA14 Δ*nahK* were used to grow 100 mL cultures of each strain in fresh LB broth to an OD 600 of 1.0. The cultures were then centrifuged at 4000 x g for 10 minutes and washed with 1X PBS once. Fresh PBS was added and cultures were adjusted to have the same OD 600. Then, 10 mLs of the suspension were mixed in with the sprouts. All sprouts were covered in a suspension of bacteria, or PBS in the case of the control, and incubated at 30C for 24 hrs.

After exposure to bacteria suspensions or PBS, each set of sprouts was rinsed with sterile ddH2O. They were then transplanted to tubes with 30 mL of Murashige-Skoog agar. Each sprout was placed in a tube and covered with After planting, plants were allowed to grow for 10 days. After 10 days, each plant was examined. Dead plants were identified by their lack of stalks and visible necrosis. After recording each plants’ status, they were removed from the agar and the weight mass was recorded. Each plant was then placed on a glass sheet and scanned on an Epson Perfection V600 scanner. Each plants’ longest root tendril and stalk length was recorded.

### Quantitative PCR (qPCR)

PA14 wildtype and *ΔnahK* were grown in 5mL succinate media for seven hours at 37°C with agitation. Total cellular RNA from 5mL cultures was isolated using the RNeasy Mini Kit (Qiagen). The yield and purity of the RNA was evaluated by Nanodrop and 1% agarose gel. cDNA was synthesized from 1 μg of RNA using the RevertAid First Strand cDNA Synthesis Kit (Thermo Scientific). The 10 μL qPCR reaction included 0.3 μM of forward and reverse primer described in Supplemental Table 1, equal amounts of cDNA and 5 μL of SYBR green master mix (Thermo Scientific). qPCR was performed on a Lightcycler 480. Cycling parameters were 95°C for 10 min, then 40 cycles of 95°C for 15 s and 60°C for 60s. *GyrA* was used as a reference gene and relative expression was determined using a standard curve (Alqarni et. al. 2016).

### LCMS

Strains were grown for 24 hours in 100mL succinate media. Supernatant was collected by centrifugation, then twice filtered before LCMS was performed, as described for other bacteria elsewhere (57). LCMS was performed on a Bruker Impact II QTOF. The data was then compared to PAMDB database (http://pseudomonas.umaryland.edu) and the Bruker database (MetaboBASE), then analyzed using Metaboanalyst software (Pang, Z., Zhou, G., Ewald, J., Chang, L., Hacariz, O., Basu, N., and Xia, J. (2022) Using MetaboAnalyst 5.0 for LC-HRMS spectra processing, multi-omics integration and covariate adjustment of global metabolomics data Nature Protocols (58).

### Construction of reporter strains

500 bp was amplified from the PA14 genome using primers listed in Supplemental Table 3 and restriction cloned upstream of the coding sequence of *mScarlet* using SpeI and XhoI digest sites in the MCS of pLD3208. Plasmids were verified by sequencing. Verified plasmids were introduced into *PA14* using biparental conjugation with *E. coli* S17-1. Single recombinants were selected on M9 minimal medium agar plates (47.8 mM Na_2_HPO_4_7H_2_O, 2 mM KH_2_PO_4_, 8.6 mM NaCl, 18.6 mM NH_4_Cl, 1 mM MgSO_4_, 0.1 mM CaCl_2_, 20 mM sodium citrate dihydrate, 1.5% agar) containing 70 µg/mL gentamicin. The plasmid backbone was removed using Flp-FRT recombination using the pFLP2 plasmid (59) and selection on M9 minimal medium agar plates containing 300 µg/mL carbenicillin. pFLP2 plasmid was cured by streaking on LB agar plates without NaCl with 10% w/v sucrose. The presence of *mScarlet* in final clones was confirmed by PCR.

### Transcriptional mScarlet Fluorescence Assays

For reporter assays, strains containing mScarlet *phz1* or *phz2* reporters were grown overnight in 5mL LB, then 1:100 diluted into black walled, clear bottom 96 well plated containing 200μL succinate media. Over 24 hours, every 30 minutes the fluorescence (ex: 560nm em: 610nm) and optical density at 600nm was determined for every well on a SpectraMax iD3 plate reader. For mixing assays, strains were mixed together in a 1:1 ratio of fluorescent, mScarlet-expressing and non-fluorescent cells. Over 24 hours, every 30 minutes the fluorescence (ex: 560nm em: 610nm) and optical density at 600nm was determined for every well on a SpectraMax iD3 plate reader (8).

## Supplemental Information

**Supplemental Table 1.** Strains used in this study.

| Strains | Relevant Characteristics | Source |
| --- | --- | --- |
| ***P. aeruginosa*** |  |  |
| UCBPP- PA14 | Wildtype | Lab Collection |
| PA14 $\Delta nahK$ | PA14 deletion mutant for PA14_38970 (*nahK*). | This study |
| PA14 *Pphz1* - mScarlet | PA14 with *Pphz1*-mScarlet inserted at the attB site using pLD3208. | This study |
| PA14 *Pphz2* - mScarlet | PA14 with *Pphz2*-mScarlet inserted at the attB site using pLD3208. | This study |
| PA14 $\Delta nahK::Pphz1$ - mScarlet | PA14 deletion mutant for PA14_38970 (*nahK*) with *Pphz1*-mScarlet inserted at the attB site using pLD3208. | This study |
| PA14 $\Delta nahK::Pphz2$ - mScarlet | PA14 deletion mutant for PA14_38970 (*nahK*) with *Pphz2*-mScarlet inserted at the attB site using pLD3208. | This study |
| $\Delta pqsABC::Pphz1$ - mScarlet | PA14 deletion mutant for PA14_51430-51410 (*pqsABC*) with *Pphz1* - mScarlet inserted at the attB site using pLD3208. | This study |
| $\Delta pqsABC::Pphz2$ - mScarlet | PA14 deletion mutant for PA14_51430-51410 (*pqsABC*) with *Pphz2*-mScarlet inserted at the attB site using pLD3208. | This study |
| $\Delta pqsR::Pphz1$ - mScarlet | PA14 deletion mutant for PA14_51340 (*pqsR*) with *Pphz1*-mScarlet inserted at the attB site using pLD3208. | This study |
| $\Delta pqsR::Pphz2$ - mScarlet | PA14 deletion mutant for PA14_51340 (*pqsR*) with *Pphz2*-mScarlet inserted at | This study |
|  | the attB site using pLD3208. |  |
| ***Escherichia coli*** |  |  |
| DH5α | *fhuA2 lac(del)U169 phoA*<br>*glnV44 Φ80' lacZ(del)M15*<br>*gyrA96 recA1 relA1 endA1*<br>*thi-1 hsdR17* | Lab Collection |
| WM3064 | *thrB1004 pro thi rpsL hsdS*<br>*lacZΔM15 RP4-1360*<br>*Δ(araBAD)567*<br>*ΔdapA1341::[erm pir]* | Lab Collection |
| S17-1 | *StrR, TpR, F-RP4-2-Tc::Mu*<br>*aphA::Tn7 recA λpir lysogen* | (60) |
| OP50 | Wildtype | Generous donation from the<br>Dr. David Q. Matus lab at<br>Stony Brook University. |
| ***Caenorhabditis elegans*** |  |  |
| N2 | Wildtype | Generous donation from the<br>Dr. David Q. Matus lab at<br>Stony Brook University. |
| ***Saccharomyces cerevisiae*** |  |  |
| InvSc1 | *MATa/MATα leu2/leu2 trp1-*<br>*289/trp1-289 ura3-52/ ura3-*<br>*52 his3-Δ1/his3-Δ1* | Invitrogen |

**Supplemental Table 2.**
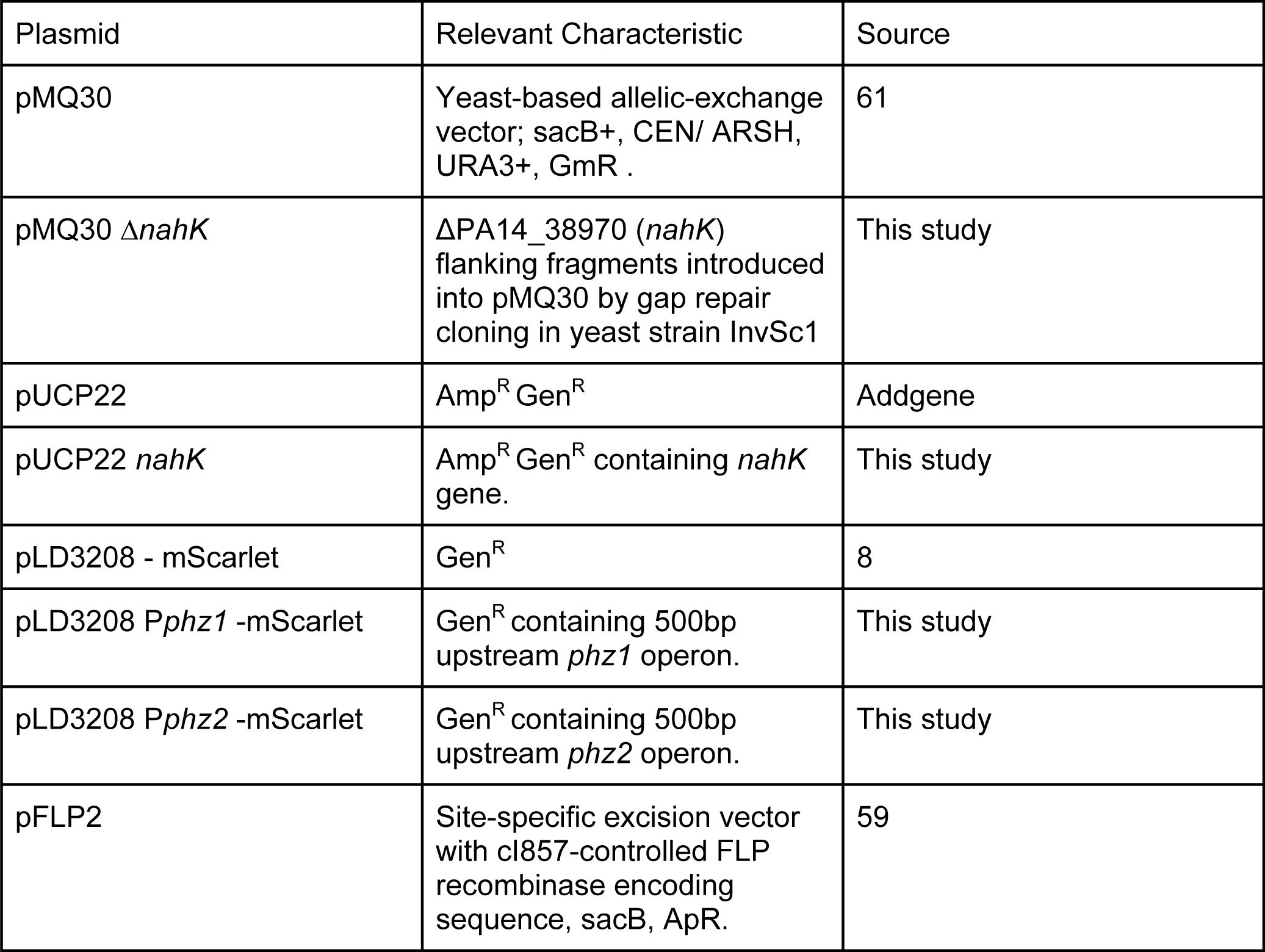
Plasmids used in this study.

**Supplemental Table 3.**
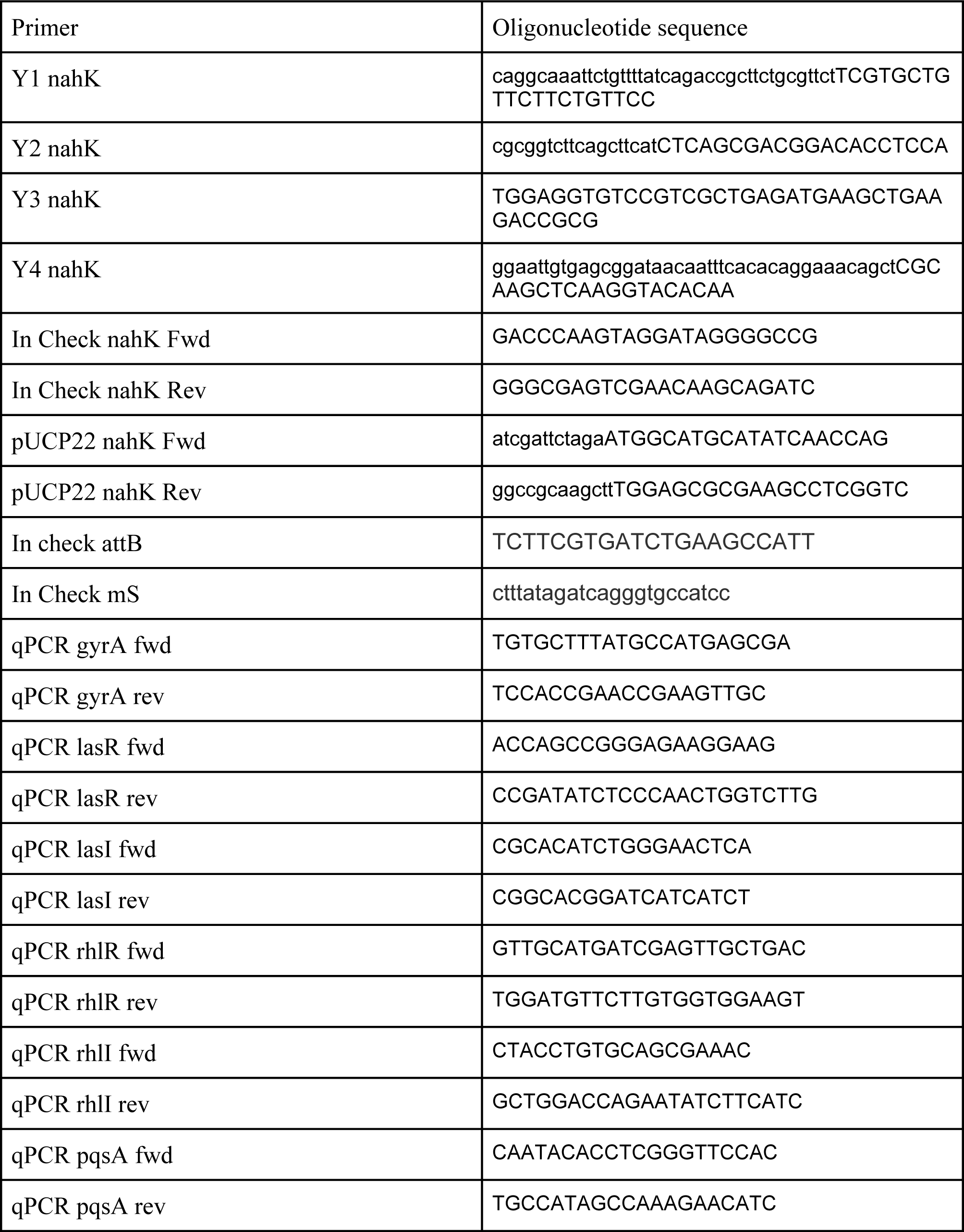
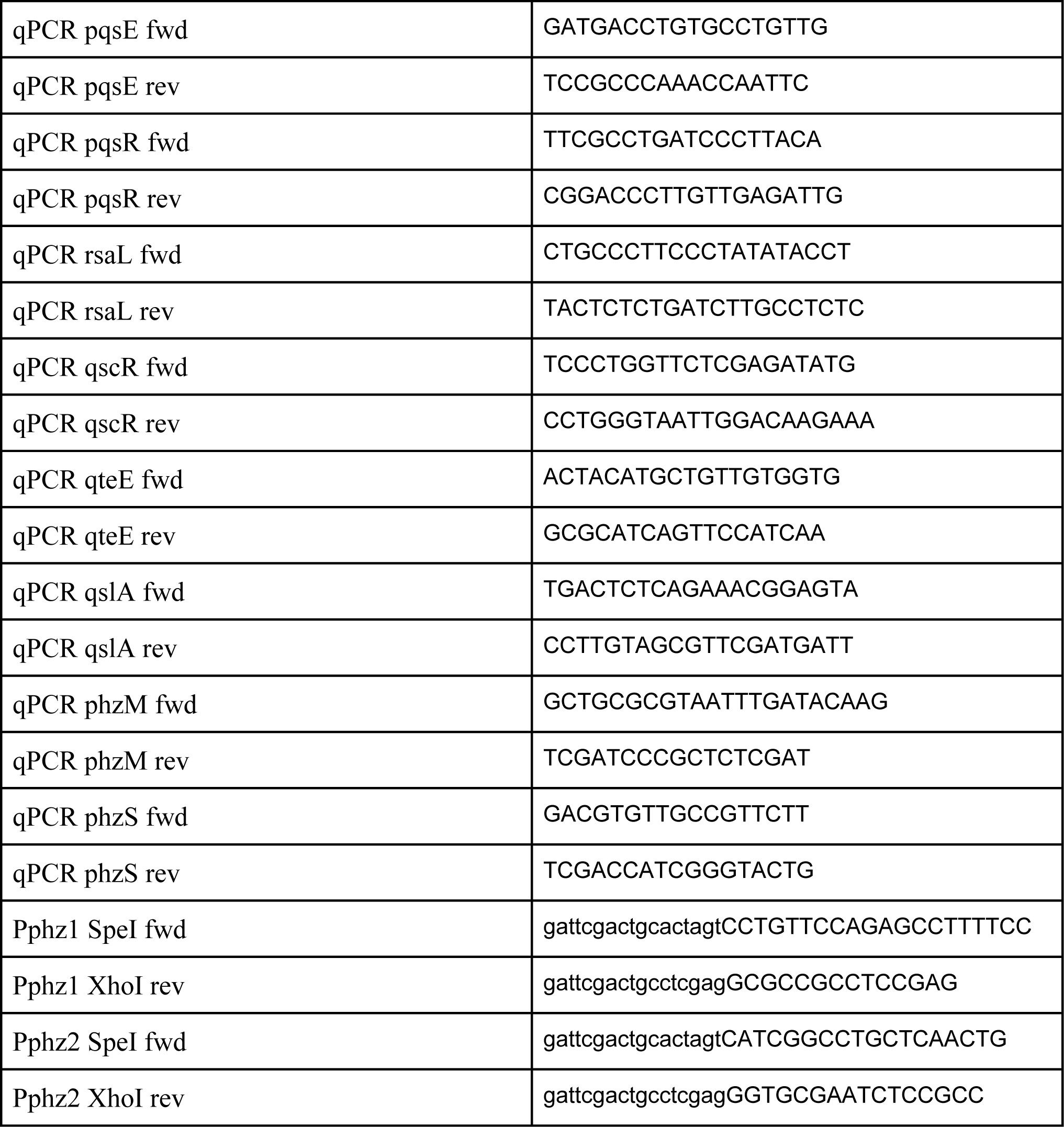
Oligonucleotides used in this study.

**Supplemental Table 4. positive & negative mode full data (ref supp fig 2)**

**Supplemental Figure 1.**
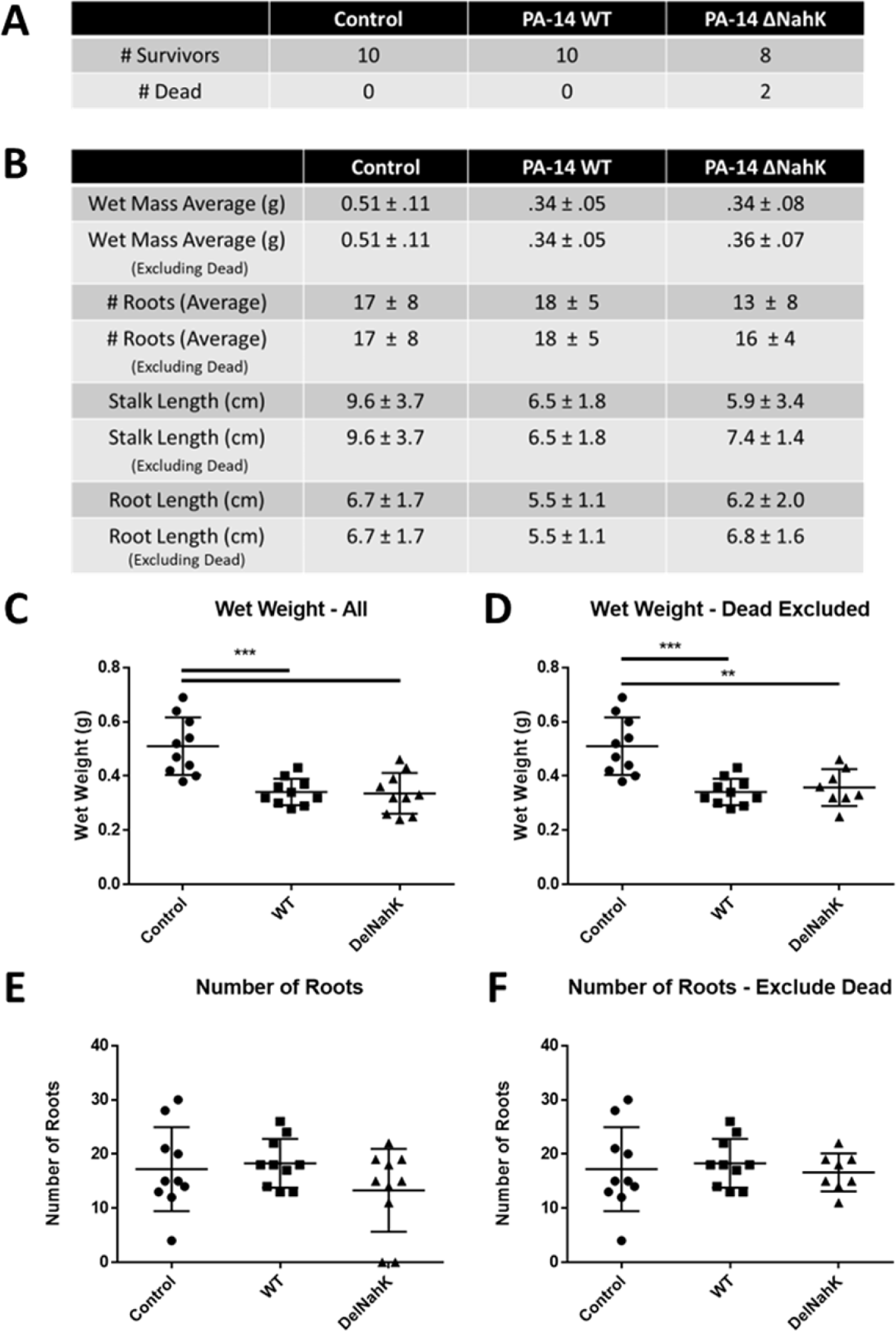
Mung bean infection model shows *ΔnahK* is more virulent than wildtype. **(A)** *ΔnahK* kills more mung bean sprouts compared to wildtype and control. **(B)** Metrics for mung bean sprout infection include wet weight, number of roots, stalk length and root length. **(C)** Wet weight of all mung beans after exposure. **(D)** Wet weight of all live mung beans after exposure. **(E)** Number of roots of all mung beans after exposure. **(F)** Number of roots of all live mung beans after exposure. ** P ≤ 0.01, *** P ≤ 0.001 one-way ANOVA with Tukey’s Multiple Comparisons.

**Supplemental Figure 2.**
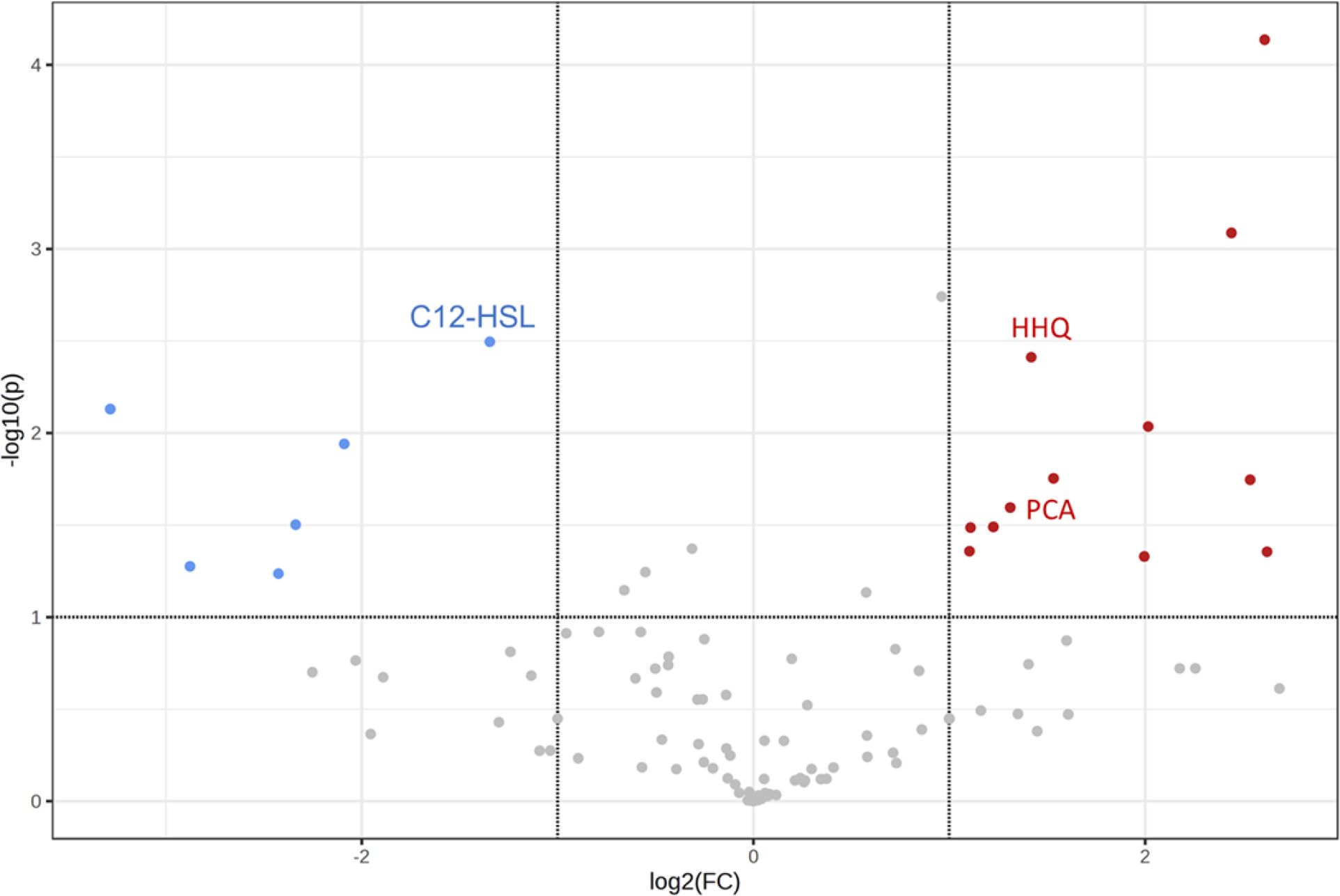
PQS precursors are overexpressed in *ΔnahK* supernatant compared to wildtype, while C12-HSL is reduced in negative mode LCMS. Values are a ratio of prevalence in *ΔnahK*:wildtype in negative mode LCMS. Bars above and below the threshold represent up- and downregulation, respectively. *p* values were calculated using unpaired, two-tailed *t-*tests unpaired, two-tailed *t-*tests (Supplemental Table 4).

**Supplemental Figure 3.**
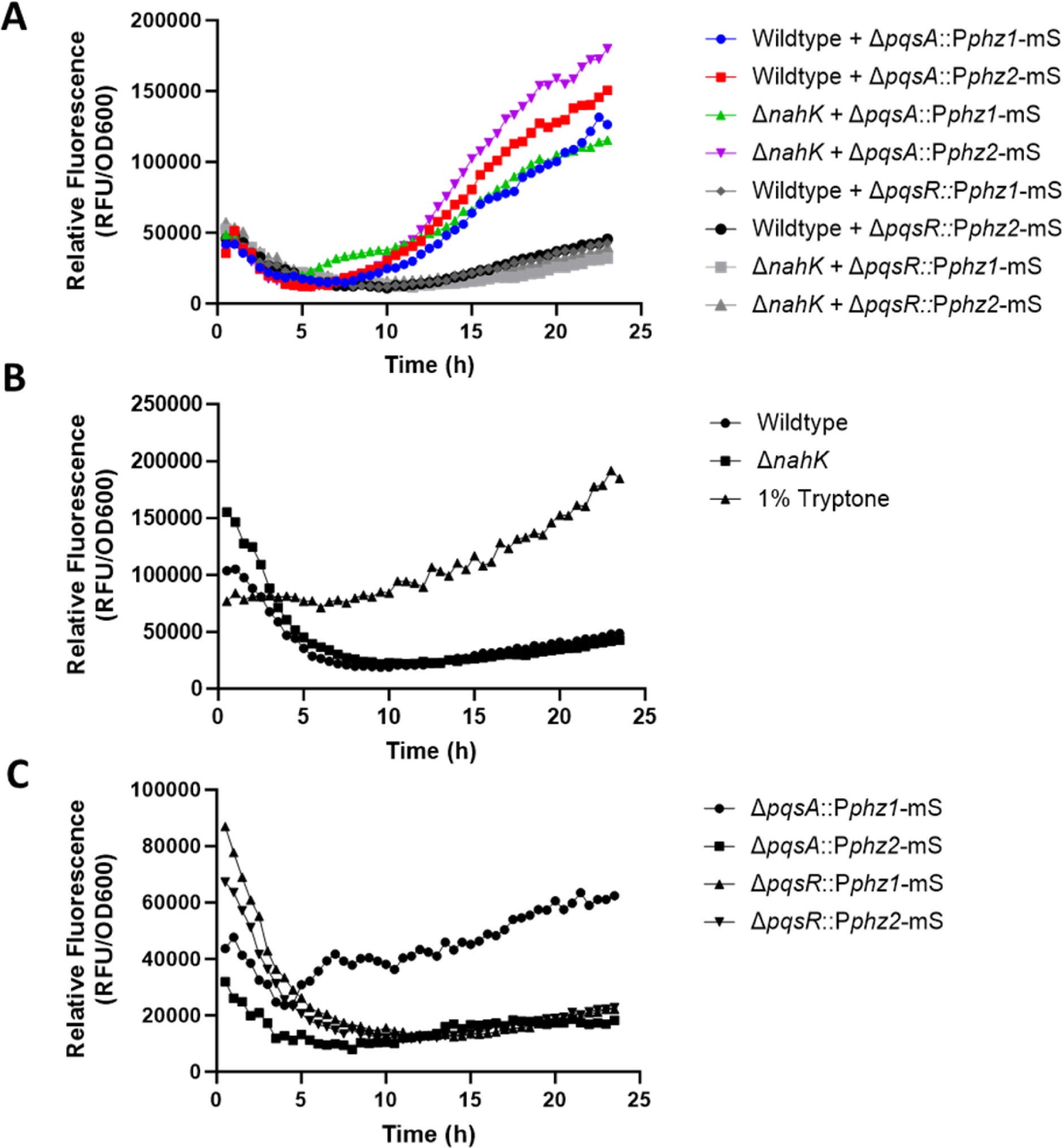
Background fluorescence from non-fluorescent and non-induced strains. Expression of mScarlet by the indicated genotypes in shaken 1% tryptone liquid cultures grown at 25 °C. n = 3 biological replicates, each with three technical replicates. Data presented as mean values. **(A)** Co-cultured wildtype with ΔpqsR::Pphz-mS strains. **(B)** Background fluorescence of donor strains and media only. **(C)** Background fluorescence of recipient strains uninduced.

